# Complementing machine learning-based structure predictions with native mass spectrometry

**DOI:** 10.1101/2022.03.17.484776

**Authors:** Timothy M. Allison, Matteo T. Degiacomi, Erik G. Marklund, Luca Jovine, Arne Elofsson, Justin L. P. Benesch, Michael Landreh

## Abstract

The advent of machine learning-based structure prediction algorithms such as AlphaFold2 (AF2) has moved the generation of accurate structural models for the entire cellular protein machinery into the reach of the scientific community. However, structure predictions of protein complexes are based on user-provided input and may therefore require experimental validation. Mass spectrometry (MS) is a versatile, time-effective tool that provides information on post-translational modifications, ligand interactions, conformational changes, and higher-order oligomerization. Using three protein systems, we show that native MS experiments can uncover structural features of ligand interactions, homology models, and point mutations, that are undetectable by AF2 alone. We conclude that machine learning can be complemented with MS to yield more accurate structural models on the small and the large scale.

## Introduction

Machine learning (ML)-based algorithms have been hailed as the solution to the protein structure prediction problem and are already being used to predict structures across entire proteomes ^1^. For example, using protein sequence data as the only user input, AF2 ^2^ can generate models of ordered, monomeric proteins that rival in quality experimentally derived structures ^3^, which can be assembled into complexes using AF2 Multimer ^4^. However, it is important to remember that the models are generated according to user-provided input. For example, AF2 Multimer does not suggest an oligomeric state; instead, the stoichiometry for the model has to be specified along with the sequences of the components. Moreover, AF2 may propose seemingly plausible models for a protein interaction even if this is not biologically relevant, for example because the proteins are in different cellular compartments. Furthermore, the use of AF2 to predict interactions that involve dynamic regions ^5^, ligand binding sites, or point mutations ^6^, all of which are major focal points of structural biology, remains challenging ^7^. In these cases, additional structural data may be required to assess the validity of the computed structures, for example from cryo-EM and X-ray crystallography. However, obtaining such data is challenging, resulting in a need for alternative strategies.

MS, with its rapidly expanding structural biology toolbox ^8^, can provide structural data that are directly complementary to ML (Figure 1A). Despite not being a stand-alone structure determination technique, MS offers a wealth of information for hybrid structural biology approaches ^8^. It has a well-developed capacity to provide proteoform primary structure information, such as post-translational modifications, via MS-sequencing. In combination with in-solution labeling methods such as hydrogen-deuterium exchange (HDX), MS can inform about local structural dynamics. Native MS, where the non-covalent interfaces in macromolecules are preserved in the experiment, is still the gold standard to determine oligomeric states, which is of particular importance when building models of protein complexes. Crosslinking and ion mobility (IM) measurements reveal the spatial arrangements of components in a protein complex. Unlike other biophysical methods, MS offers the crucial advantage of being able to provide structural data on the proteome scale. For example, proteome-wide crosslinking studies can help to filter biologically irrelevant interactions ^9^. Collision-cross sections (CCSs, effectively 2D-projections of the structures) can be calculated for entire model proteomes and used to filter complex architectures by IM-MS ^10^. Lastly, hybrid MS methods, such as NativeOmics, can reveal direct connections between primary and quaternary structure variations, as well as help to identify ligands or cofactors that may be structurally and functionally important ^11^.

**Figure 1.**
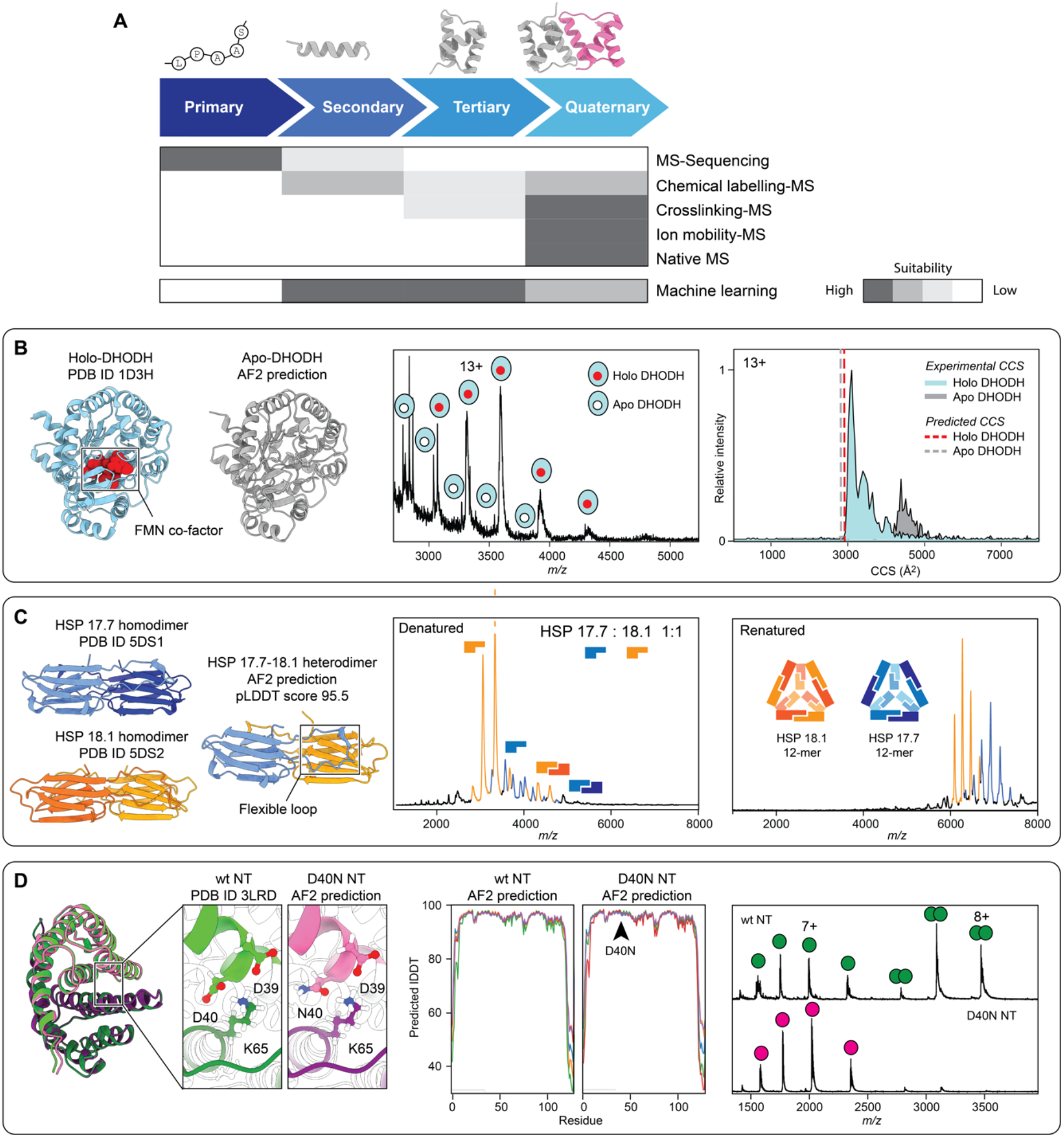
**(A)** The structural MS toolbox offers information that is directly complementary to ML-based structure prediction. MS can inform about proteoforms (MS sequencing), structural dynamics (HDX-MS), the spatial arrangements of proteins in a complex (ion mobility and crosslinking MS), and oligomeric states (native MS). **(B)** Left: Experimental and predicted structures for holo-(left) and apo-DHODH show near-identical three-dimensional folds. Middle: Native MS reveals the presence of a small population of apo protein ^14^. Right: IM-MS of the 13+ charge states of apo- and holo-DHODH shows that the protein with co-factor has a native-like CCS, whereas the protein without co-factor is unfolded. **(C)** Left: Crystal structures for the HSP17.7 and 18.1 homodimers are virtually indistinguishable from the AF2-predicted heterodimer. Native MS of a mixture of HSP17.7 and 18.1 under denaturing conditions (middle) and after refolding (right) reveal that no heterodimer formation takes place ^15^. **(D)** Left: AF2 predicts that the D40N mutant of MaSp1 NT forms a homodimer that closely resembles the dimeric structure of wt MaSp1 NT, despite showing partial loss of the D39/D40/K65 salt bridge. Middle: pLDDT plots indicate that the D40N mutation does not affect the prediction confidence for the subunits in the NT dimer. Right: Native MS analysis of both NT variants at pH 6.0 shows that the D40N mutation abolishes NT dimerization ^16^. All AF predictions were carried out using Colab Fold with AMBER step and without templates ^21^.

We therefore asked whether native MS, which is widely employed to study protein interactions, can be readily used to assess the plausibility of structural models generated by AF2. For this purpose, we selected three protein complexes whose interactions involve disordered regions, ligands, and point mutations. In all three cases, the native MS data show specific effects that are not detectable by AF2 alone, illustrating the complementarity of the two approaches.

## Results and Discussion

As a first example, we tested the ability of AF2 to predict the structure of dihydroorotate dehydrogenase (DHODH), a mitochondrial enzyme involved in uracil synthesis. Inhibition of DHODH selectively kills cancer cells, making it a prime target for the development of novel therapeutics ^12^. When using AF2 to predict the structure of the soluble domain of DHODH, the result is nearly indistinguishable from the available X-ray structures ^13^, with a root mean-square deviation (RMSD) of 0.5 Å^2^ (Figure 1B), with the exception that the predicted structure contains a central cavity which in the experimental structures is occupied by the cofactor flavin mononucleotide (FMN). We have previously used native MS to assess the relationship between ligand binding and folding of DHODH ^14^ and found that the protein exists mostly in the holo-form. We also detected a small apo population with higher charge states, indicating unfolding in solution. Indeed, IM-MS revealed that FMN-bound protein adopts a compact conformation, whereas the FMN-free protein is largely unfolded, as evident from the CCS distributions of the 13+ charge state of both populations (Figure 1B) ^14^. When we computed the CCSs of the experimental and the predicted structures, we found them to be virtually identical (Figure 1B). Taken together, we find that AF2 predicts the fold of the holo-form of DHODH even without the co-factor, whereas native MS shows that the protein cannot adopt, or maintain, the correct conformation in the absence of FMN. This discrepancy could arise from co-factor-bound proteins being part of the AF2 training set, yet the co-factors themselves are not considered in the prediction. Although alternative computational tools may be used to incorporate ligands in AF2 models, the connection between binding and folding is not considered in the predictions. As shown for DHODH, native MS can inform about the role of the co-factor in promoting the correct fold of DHODH, a role that is not evident from the ML-based prediction alone.

Next, we asked whether native MS and AF2 could capture the effect of a flexible segment on the formation of a protein complex. For this purpose, we turned to the homologous small heat shock proteins 17.7 and 18.1 from *Pisum sativum*. Both form highly similar homodimers via a conserved dimerization interface and swapping of a flexible loop, which are correctly predicted by AF2 (Figure 1C) ^15^. Using AF2, we could also predict the HSP 17.7-18.1 heterodimer with a per-residue confidence (pLDDT) score equal to those of the homodimers, and an RMSD of 0.73 Å^2^ and 0.66 Å^2^ for the 17.7 and 18.1 heterodimer, respectively. However, upon refolding a mixture of denatured HSP 17.7 and 18.1, native MS revealed homodimer formation and assembly into dodecamers, and despite no direct steric hindrance, heterodimerization is practically impossible (Figure 1C). This preference is due to an inability of the heterodimer to bind the flexible loops due to differences in non-interfacial residues, which provides a penalty for hetero-oligomerization ^15^. Such a preference of homo-over hetero-oligomerization is likely a wide-spread phenomenon ^15^. However, as it is mediated by a flexible region outside of the well-defined dimerization surface, it has no significant impact on the confidence of the AF2 model, but can be readily detected by MS.

Lastly, we investigated the ability of MS and AF2 to capture the impact of point mutations on protein complex formation. Mutations that do not introduce significant steric hindrance yield near-identical AF2 structures ^6^ that nonetheless show measurable differences in stability ^7^. However, it is unclear to what extent AF2 can inform about the effect of mutations on protein interactions. We chose the N-terminal domain (NT) of the spider silk protein Major ampullate Spidroin 1 (MaSp1) from *Euprosthenops australis*, which is monomeric above, and dimeric below, pH 6.5 ^16,17^. This pH sensitivity is in part due to a conserved salt bridge between D39/D40 and K65 on the opposing subunit ^18,19^. We used AF2 to predict the structure of the dimeric wild-type protein, as well as a point mutant with a weakened salt bridge, D40N (Figure 1 C). Importantly, AF2 does not explicitly address the protonation state of ionizable residues, but may indirectly reflect the interactions observed under the solution conditions used to solve the structures in the training set. Comparison of the pLDDT scores of the top five models for each variant showed no discernable differences (Figure 1D) with an RMSD of 0.2 Å^2^, indicating highly similar structures. Native MS analysis of both proteins at pH 6.0, on the other hand, showed that the D40N mutation abolished dimerization nearly completely (Figure 1D)^16^. In summary, mutating aspartate 40 to asparagine does not introduce structural changes or steric clashes and does not appear to have notable consequences for the F2 model of the dimer. The impact of losing this salt bridge on dimer formation therefore requires experimental validation, such as through native MS analysis.

## Conclusions

Here, we examined the ability of MS to provide complementary information to ML-based structure predictions of protein complexes. While AF2 predictions are generally of very high accuracy, they do not specifically address the influence of bound ligands, flexible regions, and point mutations on protein interactions. Native MS, on the other hand, does not provide structural details, but can capture a wide range of protein interactions with a single measurement. Of particular importance for structure prediction is the ability of MS to provide accurate information on protein oligomeric states. While MS is unrivalled in the detail of the mass measurements it provides, reliable mass measurement of multimeric stoichiometries can be obtained from various alternative techniques, opening up even more ways to complement ML predictions. Going forward, MS should be combined with ML either by defining the modelling question *a priori* using MS data (MS/AI), or by using MS data to identify a likely model *a posteriori* (AI/MS). We anticipate that whole-proteome structural MS data, and even mass measurements in physiological solutions, such as analytical ultracentrifugation and small-angle X-ray scattering, but also new methods like mass photometry ^20^, could be incorporated into large-scale ML predictions, for example in the form of constraints, to generate accurate structural maps of the entire cellular environment.

